# Stress-induced Rab11a-exosomes induce AREG-mediated cetuximab resistance in colorectal cancer

**DOI:** 10.1101/2023.12.19.572341

**Authors:** John D. Mason, Ewan Marks, Shih-Jung Fan, Kristie McCormick, Clive Wilson, Adrian L. Harris, Freddie C. Hamdy, Chris Cunningham, Deborah C. I. Goberdhan

## Abstract

Exosomes are secreted vesicles made intracellularly in the endosomal system. We have previously shown that exosomes are not only made in late endosomes, but also in recycling endosomes marked by the monomeric G-protein Rab11a. These vesicles, termed Rab11a-exosomes, are preferentially secreted under nutrient stress from several cancer cell types, including HCT116 colorectal cancer (CRC) cells. HCT116 Rab11a-exosomes have particularly potent signalling activities, some mediated by the Epidermal Growth Factor Receptor (EGFR) ligand, Amphiregulin (AREG). Mutant activating forms of KRAS, a downstream target of EGFR, are often found in advanced CRC. When absent, monoclonal antibodies, such as cetuximab, which target the EGFR and block the effects of EGFR ligands, such as AREG, can be administered. Patients, however, inevitably develop resistance to cetuximab, either by acquiring KRAS mutations or via non-genetic microenvironmental changes. Here we show that nutrient stress in several CRC cell lines causes the release of AREG-carrying Rab11a-exosomes. We demonstrate that while soluble AREG has no effect, much lower levels of AREG bound to Rab11a-exosomes from cetuximab-resistant KRAS-mutant HCT116 cells, can suppress the effects of cetuximab on KRAS-wild type Caco-2 CRC cells. Using neutralising anti-AREG antibodies and an intracellular EGFR kinase inhibitor, we show that this effect is mediated via AREG activation of EGFR, and not transfer of activated KRAS. Therefore, presentation of AREG on Rab11a-exosomes affects its ability to compete with cetuximab. We propose that this Rab11a-exosome-mediated mechanism contributes to the establishment of resistance in cetuximab-sensitive cells and may explain why in cetuximab-resistant tumours only some cells carry mutant KRAS.

**Graphical Abstract:** 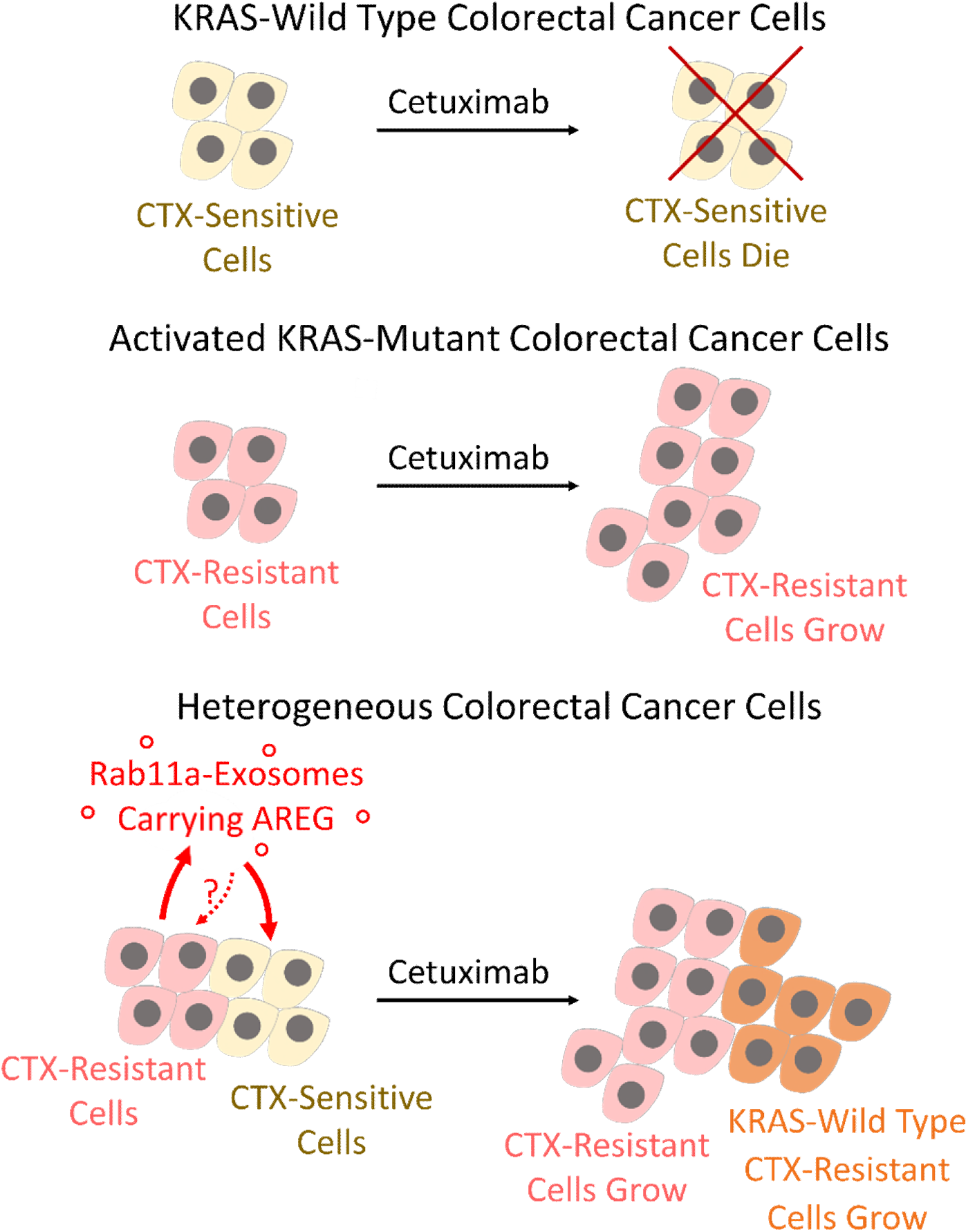

This study highlights a clinically relevant mechanism in which stress-induced Rab11a-exosomes carrying the EGFR ligand, Amphiregulin (AREG) transfer drug resistance between genetically distinct colorectal cancer cells. Resistance to cetuximab, an anti-EGFR therapy, can be passed via Rab11a-exosomes from drug-resistant KRAS-mutant cells to previously drug-responsive KRAS-wild type cells. Unlike soluble AREG, Rab11a-exosome-associated AREG competes with cetuximab to activate EGFR signalling and promote EGFR-dependent outcomes, such as growth. This mechanism may support the co-operative evolution of clonal heterogeneity during tumour progression.

## INTRODUCTION

Extracellular vesicles (EVs) are membrane-bound structures that appear to be secreted by all cell types (van Niel et al., 2022), and which deliver cargos to recipient cells that play important roles in both physiology and pathology (Yates et al., 2022; Cheng and Hill., 2022; Lucotti et al., 2022). EVs can be produced by outward protrusion and detachment from the plasma membrane, forming so-called ectosomes, or by inward budding of endosomal membranes to produce vesicles, which are secreted as exosomes, when these endosomes fuse with the plasma membrane (Dixson et al., 2023).

EVs can be separated and concentrated from conditioned medium or biofluids by methodologies based on physical properties, such as size and density, but the resulting preparations contain heterogenous mixtures of exosomes and ectosomes, as well as aggregated proteins (Jeppesen et al., 2019). For example, small EVs (sEVs) separated and concentrated by differential centrifugation (dUC) or size-exclusion chromatography (SEC) typically contain exosomes, which have a size range of 30-150 nm in diameter, but also include ectosomes of similar size. Consequently, although sEV preparations can be shown to have functional activities in target cells, it is difficult to determine whether they are mediated by the mixed preparation or a specific EV subtype with specialised activities.

Microenvironmental stress suppresses signalling by nutrient-sensitive mechanistic Target of Rapamycin Complex 1 (mTORC1; reviewed in Goberdhan et al., 2016). We have identified a specific subtype of exosome that is preferentially secreted under such stress conditions by colorectal, prostate and cervical cancer cell lines. This stress-induced change is characterised by a switch in exosome biogenesis and secretion, involving reduction in classical CD63-positive exosomes generated in late endosomal compartments and an increase in those formed in Rab11a-labelled recycling endosomes, termed Rab11a-exosomes (Fan et al., 2020). These exosomes are also made under normal physiological conditions, and in some cases constitute a key high proportion of secreted EVs, for example, in the prostate-like secondary cells of the fruit fly, *Drosophila melanogaster*, where they mediate reproductive signalling between males and females (Corrigan et al., 2014; reviewed in Wilson et al., 2017).

One major exosome biogenesis mechanism involves the endosomal complexes required for transport (ESCRT) proteins, which load exosome cargos, distort the endosomal compartment membrane during inward budding and ultimately pinch off intraluminal vesicles (ILVs; Vietri et al., 2019). We have recently shown that Rab11a-, but not late endosomal, exosome formation can be blocked by knockdown of several accessory ESCRT-III proteins (Marie et al., 2023). These experiments have revealed that Rab11a-exosomes have unique activities in physiology and disease, for example, promoting growth in the colorectal cancer (CRC) HCT116 cell line via a membrane-bound form of Amphiregulin (AREG), the ligand for the EGF receptor (EGFR, otherwise known as human EGFR-related 1, HER1) ligand (Marie et al., 2023). This exosome-associated AREG activates the EGFR at concentrations much lower than soluble AREG (Fan et al., 2020).

In the UK, colorectal cancer (CRC) is the fourth most common cancer, with an incidence of approximately 42,000 new cases and 16,000 mortalities per year (Cancer Research UK 2023). Survival is closely related to disease stage at diagnosis, ranging from 95% five-year survival for stage I to below 10% for stage IV. Surgical resection with or without neoadjuvant or adjuvant chemotherapy (or chemoradiotherapy) forms the basis of treatment with curative intent.

One important form of chemotherapy involves cetuximab, and related chimeric monoclonal antibodies, which target the EGFR and block its signalling activity (Fornasier et al., 2018). Approximately 40% of CRC patients carry KRAS mutations, and 3% have NRAS mutations, rendering anti-EGFR therapies ineffective (Bando et al., 2023). Cetuximab can, however, be used in combination treatment of metastatic CRC in patients lacking such downstream mutations, but even in these cases, it is not always effective (Allegra et al., 2016). Eventually all patients develop cetuximab resistance. In some cases, the clonal amplification of rare cells carrying activating KRAS mutations may be responsible or *de novo* KRAS mutations may emerge (Misale et al., 2012). In others, non-genetic changes in the tumour microenvironment and intercellular communication are involved (Woolston et al., 2019). Indeed, even when KRAS-mutant cells are present, they often represent a small proportion of the tumour (Misale et al., 2012). It has been proposed that they may release increased levels of EGFR ligands, such as AREG and TGF-α, which promote resistance in neighbouring KRAS-wild type cells (Hobor et al., 2014). There has only been limited progress in developing alternative drugs that can improve the survival of patients with advanced CRC (Bando et al., 2023), so a better understanding of these complex resistance mechanisms to improve patient outcome is an area of particular interest.

Here we investigate stress-induced Rab11a-exosome secretion from several different CRC cell lines and test whether this exosome sub-type can mediate cetuximab resistance. We show that membrane-bound AREG located on Rab11a-exosomes from nutrient-stressed, KRAS-mutant HCT116 cells can compete with cetuximab and induce resistance in cells that do not carry KRAS mutations, even when much higher concentrations of soluble AREG have no effect. Our work suggests a novel mechanism by which resistance to anti-EGFR therapy can be established, producing the heterogeneous clonal populations that are frequently observed in drug-resistant patients.

## RESULTS AND DISCUSSION

### Rab11a-exosome secretion from a range of CRC cell lines is increased in response to mTORC1 inhibition

A panel of four cell lines (Fig. 1a) representing some of the molecular, cellular and genetic heterogeneity observed in CRC, namely microsatellite instability (MSI)/stability (MSS) (Flecchia et al., 2022); consensus molecular subtype (CMS) (Guinney et al., 2015; Sveen et al., 2018); proteomic signature (Wang et al., 2017); and oncogene/tumour suppressor status (Ahmed et al., 2013), was investigated to determine whether these cells all secrete more Rab11a-exosomes under nutrient stress. Cells were cultured in serum-free conditions, but the medium was supplemented with insulin, selenium and transferrin (ITS) to maintain growth factor signalling. Size-exclusion chromatography (SEC) was employed to separate the secreted sEVs.

**Fig 1.**
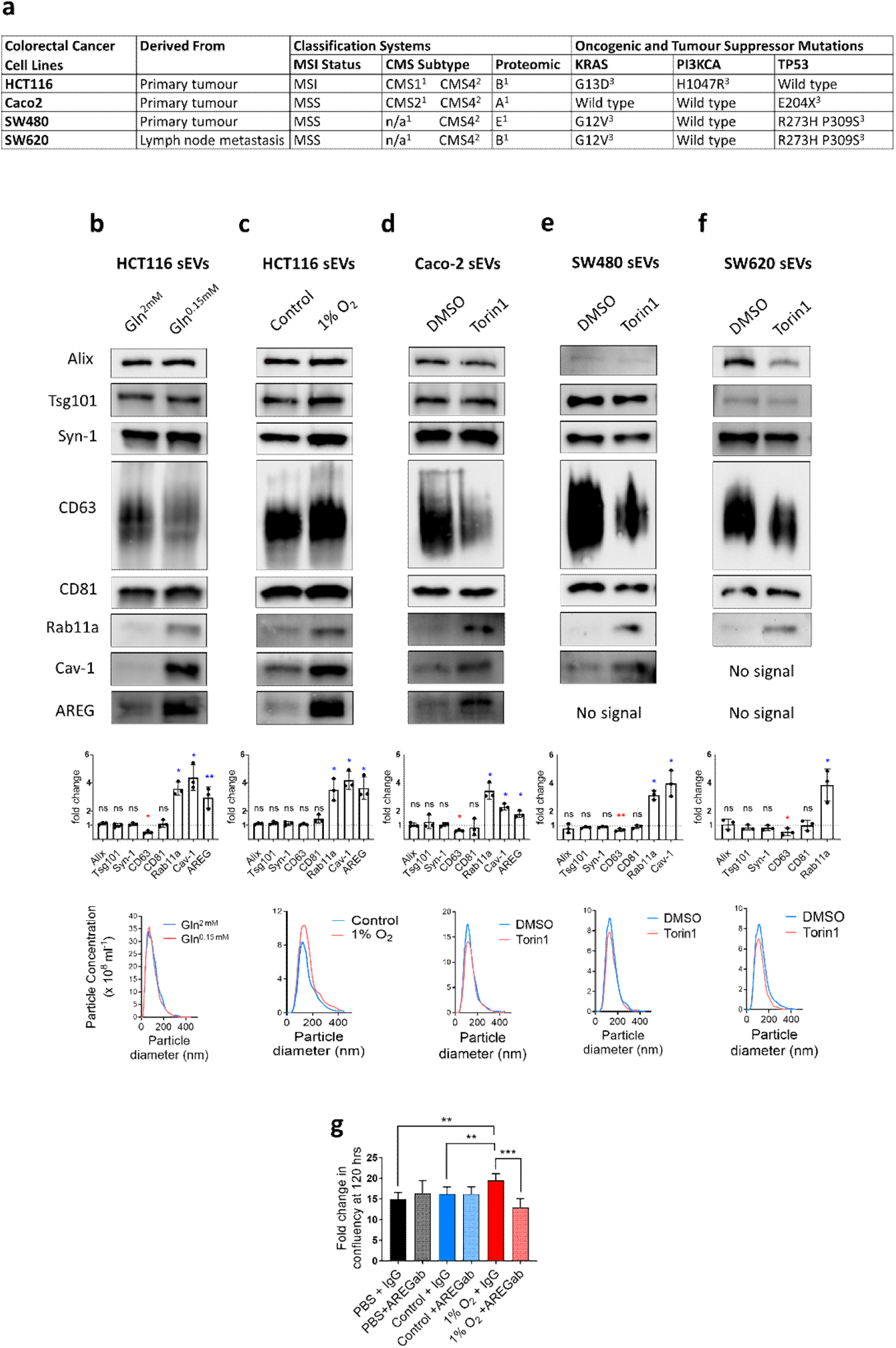
Inhibition of mTORC1 induces secretion of Rab11a-exosomes in multiple CRC cell lines. (a) Classification of the four CRC cell lines investigated based on molecular pathways and mutational status in critical oncogenes and tumour suppressor genes. (b) Western blot analysis of sEVs isolated from HCT116 cells cultured in glutamine-depleted (Gln^0.15 mM^) versus control glutamine-replete (Gln^2 mM^) conditions. Note characteristic increase in Rab11a, AREG and Cav-1 levels and the reduction in CD63 that are consistently observed (see histogram, where changes in protein levels, measured by densitometry and normalised to protein levels in secreting cell lysates, are plotted for three independent experiments; mean ± SD). NTA analysis reveals that the total number and size distribution of secreted particles is unchanged by this treatment. (c) Western blot of sEV proteins isolated from HCT116 cells cultured in hypoxic (1% O_2_) versus normoxic conditions also reveals an increase in markers of Rab11a-exosomes (Rab11a and AREG) and other stress-induced sEVs (Cav-1). (d-f) Western blots of sEV proteins from Caco-2 (d), SW480 (e) and SW620 (f) cells treated with the ATP-competitive mTORC1 inhibitor, Torin1 (100 nM for SW480 cells and 150 nM for Caco-2 and SW620 cells), versus treatment with vehicle alone (DMSO) also reveals a switch to secretion of increased Rab11a and AREG, and a reduction in CD63. NTA reveals that all sEV preparation have similar size distribution, but particle numbers are about 25% reduced. (g) Graph showing levels of HCT116 growth after 120 h, measured as fold change in confluency, following addition of PBS, control HCT116 sEVs, or hypoxia-induced HCT116 sEVs in the presence or absence of pre-incubation with neutralising anti-AREG antibodies. Note that only sEVs isolated under hypoxic conditions enhance growth and this is AREG-dependent. *P<0.05. Red and blue asterisks denote reduction and increase in protein levels respectively in Rab11a-exosome-enriched sEVs. Abbreviations used: MSI (microsatellite instability), MSS (microsatellite stability), CMS (consensus molecular subtype), n/a (not applicable). KRAS activating mutations: G13D or G12V. PI3K activating mutations: E545K D549N or H1047R. TP53 inactivating mutations: R273H P309S, or S241F, or E204X. Superscripts ^1,2,3^ refer to ^1^Wang *et al*., 2017; ^2^Sveen *et al*., 2018; ^3^Ahmed *et al*., 2013.

In HCT116 cells, downregulation of the mTORC1 signalling pathway in response to glutamine depletion, confirmed by western blot analysis of mTORC1 targets, 4E-BP1 and S6 (Fig. S1a, b), led to a switch to increased secretion of Rab11a-exosomes carrying membrane-bound AREG (Fig. 1b; cf. Fan et al., 2020; Marie et al., 2023). As previously shown, there was also an increase in sEV-associated levels of the scaffolding protein Caveolin-1 (Cav-1). Experiments using immuno-affinity-separation of sEVs and selective inhibition of Rab11a-exosome secretion suggest that Cav-1, unlike AREG, is associated with alternative stress-induced vesicles that co-separate with Rab11a-exosomes (Fan et al., 2020; Marie et al., 2023). Consistent with previous studies, glutamine depletion did not affect the expression levels of other exosome and EV proteins in cells, except for CD63, which was reduced under stress conditions in both sEVs (Fig. 1b) and cells (Fig. S1b). Furthermore, there was no detectable change in the total number or the size range of these sEVs, as determined by Nanoparticle Tracking Analysis (NTA; Fig. 1b). Analysis of SEC-separated sEVs by transmission electron microscopy (TEM) revealed the standard cup-like morphology typically associated with sEV preparations (Fig. S1g).

We also found that hypoxia, a common microenvironmental stress in fast-growing tumours, previously shown to alter sEV cargos in glioblastoma cells (Kurcharzewska et al., 2013), inhibited mTORC1 (Fig. S1c) and increased Rab11a-exosome secretion (Fig. 1c). In contrast to glutamine depletion, the levels of sEV-associated CD63 were not reduced under hypoxia, suggesting that secretion of late endosomal exosomes was not reduced by this treatment. Hypoxia-induced vesicles preferentially promoted growth of HCT116 cells under serum-depleted conditions, an activity that could be blocked by adding neutralising anti-AREG antibodies to the sEV preparations (Fig. 1g). A similar inhibitory effect was previously observed on Rab11a-exosome-enriched sEV preparations from glutamine-depleted HCT116 cells (Fan et al., 2020) and suggests that the enhanced growth-promoting effects are mediated by membrane-associated AREG loaded on to hypoxia-induced Rab11a-exosomes.

For Caco-2, SW480 and SW620 CRC cell lines, glutamine depletion did not affect mTORC1 activity in a dose-response experiment, as determined by the phosphorylation state of downstream target 4E-BP1, and only variably reduced S6 phosphorylation (Fig. S2a-d). Each of these cell lines was, however, sensitive to the mTORC1 inhibitor Torin1 in the dose range of between 100-150 nM, which affected both S6 and 4E-BP1 (Fig. S2e-g). Under these conditions, Torin1 treatment induced an increase in sEV-associated Rab11a and Cav-1 (although not for SW620 cells, which lack Cav-1), and a decrease in the tetraspanin CD63 (Fig 1d-f), but without a change in levels of these proteins in cell lysates (except for reduced CD63 levels in Caco-2 cells; Fig. S1d-f). Only sEVs from Caco-2 cells contained detectable levels of AREG, but as with HCT116 cells, these levels were strongly increased following mTORC1 inhibition. In Caco-2 cells, however, cellular levels of AREG were not affected by this treatment. Other EV markers, such as Alix, Tsg101, Syn-1 and CD81, were unchanged in sEV preparations from all three cell types (Fig 1d-f) and also in the secreting cells (Fig S1d-f). For all three cell lines, Torin1 treatment did not significantly alter the size of vesicles in sEV preparations: Caco-2, 139 ± 58 nm diameter (control) versus 143 ± 57 nm (Torin1); SW480, 149 ± 55 nm (control) versus 150 ± 49 nm (Torin1); SW620, 111 ± 54 nm (control) versus 109 ± 54 (Torin1). sEV secretion, however, was reduced by about 25% for each of these cell lines, as determined by NTA (Fig 1d-f). EVs from each cell line produced cup-like particles typical of sEV morphology in TEM studies (Fig S1h-j).

In summary, CRC cell lines with different molecular, cellular and genetic properties can be induced to release increased levels of Rab11a-exosomes by reducing cellular mTORC1 signalling, suggesting that secretion of these alternative sEVs is a typical response to stresses that suppress this nutrient sensor. Furthermore, for CRC cells in which exosomal AREG can be detected, namely HCT116 and Caco-2, a membrane-associated form of AREG is also preferentially secreted under these stress conditions.

### HCT116- and Caco-2-derived Rab11a-exosome preparations can induce AREG-dependent growth in Caco-2 cells

Rab11a-exosome preparations isolated from HCT116 cells under glutamine-depleted conditions can selectively induce AREG-dependent growth of naïve HCT116 cells cultured in low (1%) serum (Fan et al., 2020; Marie et al., 2023). We confirmed these findings using a range of sEV concentrations (Fig. 2a, b). As previously found, sEVs isolated from glutamine-replete cells had no detectable growth-promoting activity. The Rab11a-exosome preparations stimulated growth in a dose-dependent fashion (Fig. S3a, b), again consistent with our previous observations (Fan et al., 2020).

**Fig. 2.**
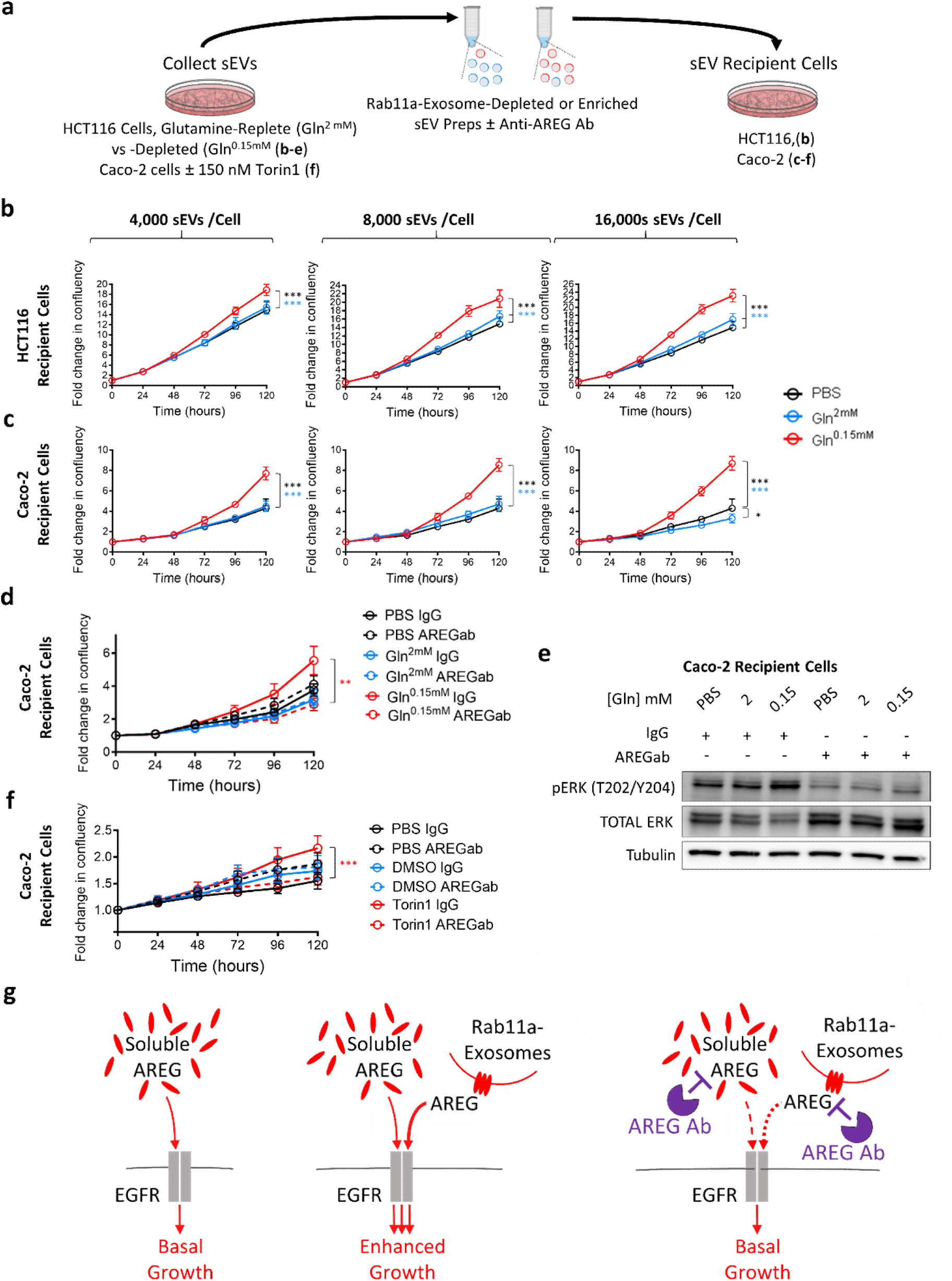
Rab11a-exosome preparations from HCT116 and Caco-2 cells enhance growth of HCT116 and Caco-2 cells in an AREG-dependent fashion. To test the effect of Rab11a-exosome-enriched sEV preparations (red lines on graphs) versus control sEV preparations (blue lines on graphs) on cell growth, sEVs were isolated either from HCT116 cultured in glutamine-depleted (Gln^0.15 mM^) versus glutamine replete (Gln^2 mM^) conditions; or from Caco-2 cells cultured in 150 nM Torin1 versus vehicle and compared to PBS addition alone (black line on graphs). They were then added to HCT116 or Caco-2 recipient cells in reduced serum conditions (1% and 0.25% FBS respectively). (a) Diagram of experimental design. (b) Compared to control sEV preparations (blue line), Rab11a-exosome-enriched sEV preparations (red line) promote the growth of HCT116 recipient cells in a dose-dependent manner (see Fig. S3b). (c) These preparations were also able to promote the growth of Caco-2 recipient cells, but even the lowest dose of sEVs appeared to produce near-maximal stimulation (see Fig. S3c). (d) The additional Caco-2 cell growth stimulated by Rab11a-exosome-enriched sEVs isolated from HCT116 cells is blocked by pre-incubation with an anti-AREG antibody, which does not significantly affect growth under other conditions. (e) The anti-AREG antibody blocks the increase in levels of activated phospho-ERK (P-ERK) produced in Caco-2 cells treated with HCT116 Rab11a-exosome preparations. (f) The additional Caco-2 cell growth stimulated by Rab11a-exosome-enriched sEVs isolated from Caco-2 cells (red solid line) is blocked by pre-incubation with an anti-AREG antibody (red dashed line), which does not significantly affect growth under other conditions. (g) Schematic showing the enhanced growth-promoting effect of Rab11a-exosomes, carrying the EGFR ligand AREG (middle), compared to free AREG in serum (left). The additional Rab11a-exosome-induced growth is blocked by addition of an anti-AREG antibody (right). Eight technical repeats were employed for each condition and each experiment was repeated three times. **P<0.01; ***P<0.001. Black, blue and red asterisks denote significantly different from PBS control, control sEVs and anti-AREG antibody treated sEVs respectively.

HCT116 cells carry activating mutations in KRAS and PI3KCA, which regulate two major signalling pathways downstream of the EGFR, making them resistant to EGFR inhibition. Nevertheless, CRC tumours are typically heterogenous, consisting of multiple genetically different clones, some of which carry wild type forms of these oncogenes, even within tumours that are resistant to EGFR-targeting chemotherapies (Misale et al., 2012). We tested whether the growth-promoting activity of HCT116 Rab11a-exosome preparations could also stimulate the growth of cells that did not carry these mutations. Indeed, these preparations selectively promoted the growth of Caco-2 cells, which lack these mutations, under reduced (0.25%) serum conditions (Fig. 2c). These cells were relatively sensitive to Rab11a-exosome preparations; adding 16 x 10^3^ versus 4 x 10^3^ sEVs per target cell produced a small, but non-significant, increase in growth (Fig. S3c). Consistent with previous experiments using HCT116 target cells, this effect was blocked by a neutralising anti-AREG antibody (Fig. 2d), which suppressed the sEV-induced activation via phosphorylation of the downstream signalling target ERK (Fig. 2e, g).

Since mTORC1-inhibited Caco-2 cells also secrete increased levels of Rab11a-exosomes and AREG, when compared to control Caco-2 cells with uninhibited mTORC1 signalling (Fig. 1d), we tested whether sEV preparations from these former cells might also stimulate Caco-2 cell growth in an AREG-dependent fashion. Indeed, Rab11a-exosome-enriched sEVs secreted from Torin1-treated Caco-2 cells produced mild growth-promoting effects on Caco-2 cells cultured in low serum and these effects were selectively inhibited following incubation with an anti-AREG antibody (Fig. 2f, g), suggesting that AREG associated with stress-induced Rab11a-exosomes from multiple CRC cell lines can promote growth.

Overall, we conclude that under stress conditions, CRC cells secrete increased levels of Rab11a-exosomes, which can stimulate the growth of their neighbours through paracrine signalling involving exosome-associated AREG.

### Rab11a-exosomes can induce resistance to cetuximab through an AREG-dependent mechanism

EGFR-targeted monoclonal antibodies, such as cetuximab, are used to treat KRAS-wild type metastatic colorectal cancer, but eventually, resistance to this therapy emerges (Sforza et al., 2016; Bhattacharya 2023). Since sEVs from CRC cells have previously been reported to induce cetuximab resistance (Zhang et al., 2017; Yang et al., 2018), we set out to test whether HCT116 sEVs, and in particular Rab11a-exosomes, might mediate such effects.

Treatment of KRAS-mutant HCT116 cells and KRAS-wild type Caco-2 cells with cetuximab under serum-replete and -depleted conditions confirmed that while there was no effect on HCT116 cell growth (Fig. S4a, b), Caco-2 cell growth was sensitive to this drug under both conditions (Fig. S4c). Furthermore, analysis of the phosphorylation state of both the EGFR and downstream ERK in these two cell lines under reduced serum conditions demonstrated that cetuximab selectively inhibited EGFR signalling in Caco-2 cells (Fig. S4d).

In cases where clones of cells carrying activating KRAS mutations emerge in tumours that are resistant to anti-EGFR therapies, it is notable that most of the cells in the tumour do not carry this mutation (Misale et al., 2012). We hypothesised that sEVs, and in particular Rab11a-exosomes, released from small numbers of cetuximab-resistant CRC cells under cellular stress, might transfer resistance to cetuximab-sensitive cells, permitting the survival and growth of these cells through paracrine signalling. We therefore undertook growth assays, incubating recipient Caco-2 cells in serum-replete and -depleted conditions with or without stress-induced sEVs from HCT116 cells, and in the presence or absence of cetuximab (Fig. 3a).

**Fig. 3.**
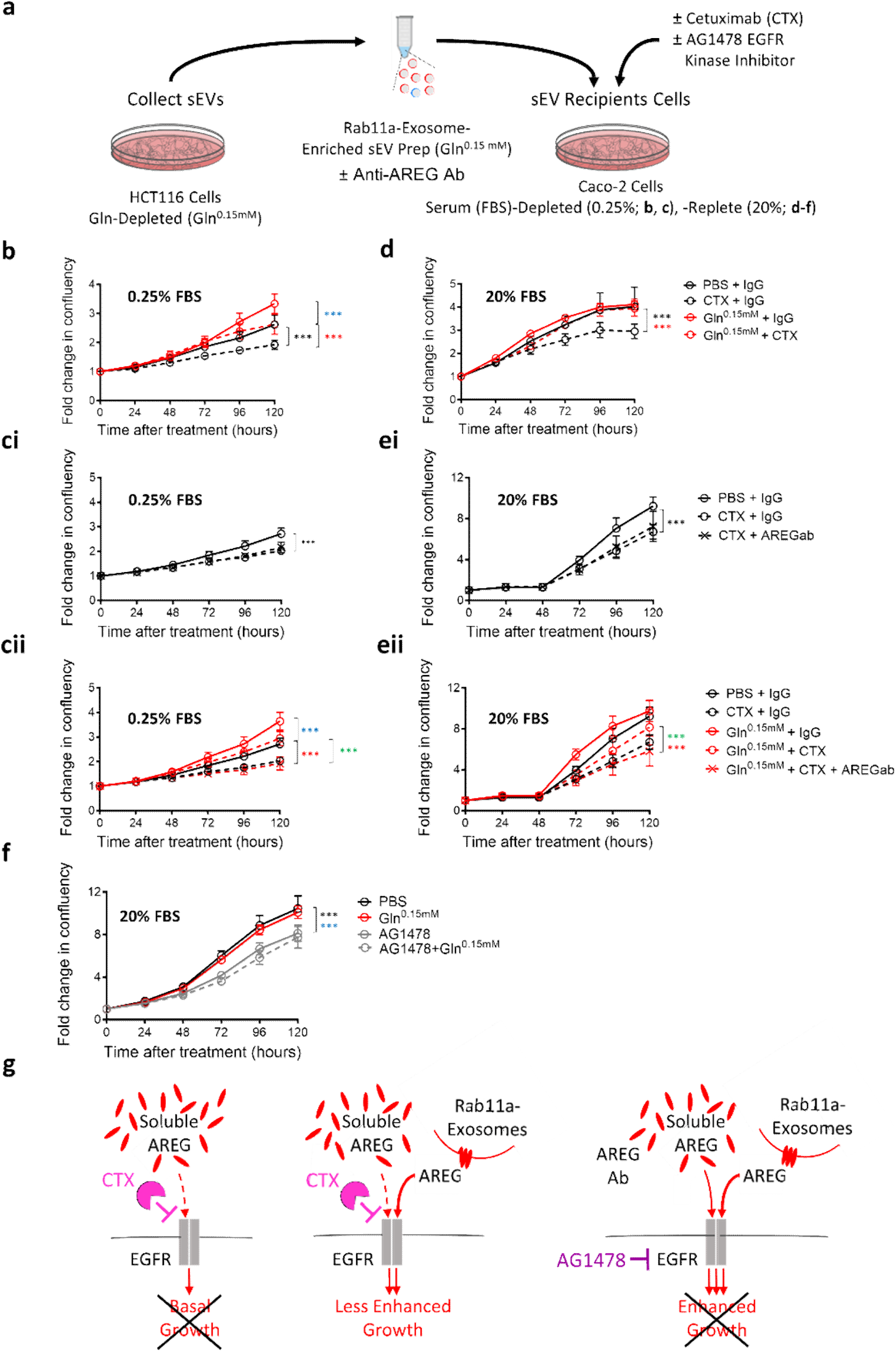
Rab11a-exosome preparations induce AREG-dependent cetuximab resistance in Caco-2 cells. (a) Rab11a-exosome-enriched sEV preparations isolated from HCT116 cells were tested for their effects on cetuximab resistance in Caco-2 cells, both in serum-depleted (0.25% FBS) and serum-replete (20% FBS) conditions, in the absence or presence of anti-AREG antibodies and an EGFR kinase inhibitor. In b-e, red and black lines on growth curves represent cells treated with Rab11a-exosome-enriched sEVs and PBS respectively, and dashed lines represent cetuximab-treated cells. (b) HCT116 Rab11a-exosome-enriched sEV preparations suppress the growth inhibitory effect of cetuximab on Caco-2 cells in serum-depleted (0.25% FBS) conditions. (ci) Addition of anti-AREG antibodies (AREG ab) to control Caco-2 cells (PBS) treated with cetuximab has no further growth inhibitory effect on cetuximab-treated cells (data points marked with crosses). (cii) In the same experiment, however, these antibodies completely suppress the growth-promoting and cetuximab-resistant effects of Rab11a-exosome preparations (red lines). Control cell growth curves with and without cetuximab (black lines) are included in this graph for comparison. (d) HCT116 Rab11a-exosome-enriched sEV preparations suppress the growth inhibitory effect of cetuximab on Caco-2 cells in serum-replete (20% FBS) conditions, where in the absence of cetuximab, they have no additional growth-promoting activity. (ei) Addition of anti-AREG antibodies to control Caco-2 cells has no further growth inhibitory effect on cetuximab-treated cells. (eii) In the same experiment, however, these antibodies completely suppress the growth-promoting and cetuximab-resistant effects of Rab11a-exosome preparations (red lines). Control cell growth curves with and without cetuximab are included in this graph for comparison (black lines). (f) The growth-promoting effects of HCT116 Rab11a-exosome preparations are completely suppressed by the EGFR kinase inhibitor AG1478. (g) Schematic models illustrating the effect of Rab11a-exosomes, carrying the EGFR ligand AREG (middle), but not AREG in serum (left), in promoting cetuximab resistance in Caco-2 CRC cells. Addition of either anti-AREG antibodies (not shown) or the EGFR kinase inhibitor AG1478 (right) blocks the cetuximab resistance induced by Rab11a-exosomes. Eight technical repeats were employed for each condition and each experiment was repeated three times. ***P<0.001. In b-e, blue asterisks denote that sEV-treated cell growth is significantly increased compared to PBS control. Red and green asterisks denote that sEV- and cetuximab-treated cell growth is significantly increased compared to PBS control cells treated with cetuximab, and sEV-/cetuximab-treated cells after sEV pre-incubation with anti-AREG antibodies respectively. In f), black (control) and blue (sEV-treated) asterisks denote significantly reduced cell growth after AG1478 treatment.

In low (0.25%) serum conditions, addition of stress-induced sEVs from glutamine-depleted HCT116 cells significantly stimulated the growth of Caco-2 cells, relative to addition of PBS alone (Fig. 3b). We have previously shown that these sEV preparations from HCT116 cells carry pg/ml concentrations of sEV-associated AREG (Fan et al., 2020). Reproducing these growth-promoting effects with recombinant AREG required ng/ml concentrations (Fig. S5a, b), confirming our previous finding that Rab11a-exosomes induce growth at much lower concentrations of AREG than soluble ligand.

Cetuximab reduced the growth of control (PBS) cells, suggesting that part of the growth observed in low serum conditions is stimulated by EGFR ligands. Cetuximab also reduced the growth induced by Rab11a-exosomes, but only to levels observed in control cells that were not treated with cetuximab (Fig. 3b). Growth was significantly enhanced compared to cells treated with PBS and cetuximab, indicating that stress-induced sEVs induce cetuximab resistance. Importantly, although adding an anti-AREG antibody had no additional growth-inhibitory effect on cells treated with PBS and cetuximab (Fig. 3ci), it further suppressed growth in the presence of HCT116 Rab11a-exosomes and cetuximab to the same levels as cetuximab-treated control cells (Fig. 3cii). This suggests that the cetuximab resistance mediated by these vesicles is AREG-dependent and therefore associated with Rab11a-exosomes.

In high (20%) serum conditions, addition of HCT116 Rab11a-exosome preparations had no significant effect on the growth of Caco-2 cells relative to PBS addition, presumably because growth was already maximally stimulated by growth factors in 20% serum (Fig. 3d). Cetuximab, however, had a completely different effect on cell growth in these two conditions. While it strongly suppressed growth in control (PBS) Caco-2 cells, it had no significant effect on growth in the presence of stress-induced HCT116 sEVs (Fig. 3d).

As in low serum conditions, addition of an anti-AREG antibody had no further growth-inhibitory effect on control (PBS) Caco-2 cells (Fig. 3ei), but it reduced the growth of Caco-2 cells in the presence of HCT116 sEVs to the levels observed in cetuximab-treated control cells (Fig. 3eii).

Taken together, these data indicate that HCT116 stress-induced sEV preparations containing Rab11a-exosomes confer cetuximab resistance to KRAS-wild type Caco-2 cells by competing with cetuximab at the EGFR (summarised in Fig. 3g). There are, however, other interpretations. In particular, AREG on Rab11a-exosomes could be acting via a receptor other than the EGFR, or the anti-AREG antibody could indirectly block another signalling activity on Rab11a-exosomes when it binds at the surface of these vesicles. We reasoned that in either of these scenarios, the HCT116 Rab11a-exosome preparations would drive cell growth, even if the EGFR was blocked in a way that did not directly compete with AREG. To test this, we added the EGFR kinase inhibitor, AG1478 (Bishop et al., 2002), to Caco-2 cells treated with PBS or HCT116 Rab11a-exosomes. In this case, growth was similarly inhibited (Fig. 3f, g), demonstrating that the effects of Rab11a-exosomes on growth are being mediated through the EGFR, even though they are cetuximab-resistant. The pan-HER receptor kinase inhibitor BMS-599626 (Wong et al., 2006) also completely blocked the growth-promoting effects of HCT116 Rab11a-exosome preparations (Fig. S5c, d), consistent with this interpretation.

The emergence of cetuximab resistance is an inevitable outcome of cetuximab treatment of KRAS-wild type tumours in metastatic CRC patients. It either involves the amplification of CRC cells carrying an activated KRAS mutation, which constitute a minority clone within a heterogeneous group of wild type KRAS cells (Misale et al., 2012), or the development of cetuximab resistance through microenvironmental changes (Woolston et al., 2019). Whatever the explanation, clonal heterogeneity and paracrine signalling appear to play a key role in this process (Chan and Buczaki, 2021). Indeed, Hodor et al. (2014) have reported an important function for EGFR ligands like AREG in mediating the communication between KRAS-mutant and KRAS-wild type cells in heterogeneous cetuximab-resistant CRC tumours.

We have shown that this resistance can be mediated by AREG-loaded, Rab11a-exosomes produced by CRC cells under nutrient or hypoxic stresses, which are commonly experienced by growing tumours. We have previously demonstrated that AREG on these vesicles can enhance CRC growth at concentrations of approximately 1 pg/ml under conditions where soluble AREG requires concentration three orders of magnitude higher (Fan et al., 2020). Here, we find that even under serum-replete conditions, where addition of Rab11a-exosomes from mutant KRAS HCT116 cells has no additional growth-promoting effect on KRAS-wild type Caco-2 cells, these exosomes, but not soluble AREG, can compete with cetuximab for EGFR binding and activation. We suggest that the mechanism by which membrane-bound AREG is presented to the EGFR, which may involve AREG clustering or association with other surface proteins, allows this ligand to circumvent the cetuximab blockade, which is normally mediated by high affinity monoclonal antibody binding. Delivery of AREG to target cells in Rab11a-exosome or soluble forms may explain why high levels of AREG have been associated both with improved and poorer outcomes in response to cetuximab treatment (Williams et al., 2023; Randon and Pietrantonio, 2023; Hong et al., 2020).

Previous studies have suggested that CRC exosomes can promote cetuximab resistance via the PTEN/Akt pathway, another target of EGFR (Zhang et al., 2018), or via the transfer of the long non-coding RNA uroepithelial carcinoma-associated-1 (UCA-1) (Yang et al., 2018). Our data suggest an alternative mechanism that may explain how KRAS-mutant cells could promote survival of KRAS-wild type cells in a heterogeneous tumour (Fig. 4). These findings suggest that treatments that either block Rab11a-exosome biogenesis or suppress the active cargos on these exosomes might act synergistically with anti-EGFR therapies to inhibit tumour growth and either slow or halt recurrence. In this regard, it is interesting to note that other forms of drug resistance have been attributed to exosome-associated ligands, such a resistance to immune checkpoint inhibitors that re-activate the patient’s anti-tumour response (Chen et al., 2018; Poggio et al., 2019). It will be interesting to explore whether stress-induced Rab11a-exosomes are involved in this resistance mechanism and whether selectively suppressing the production of these vesicles might provide a new therapeutic strategy.

**Fig. 4.**
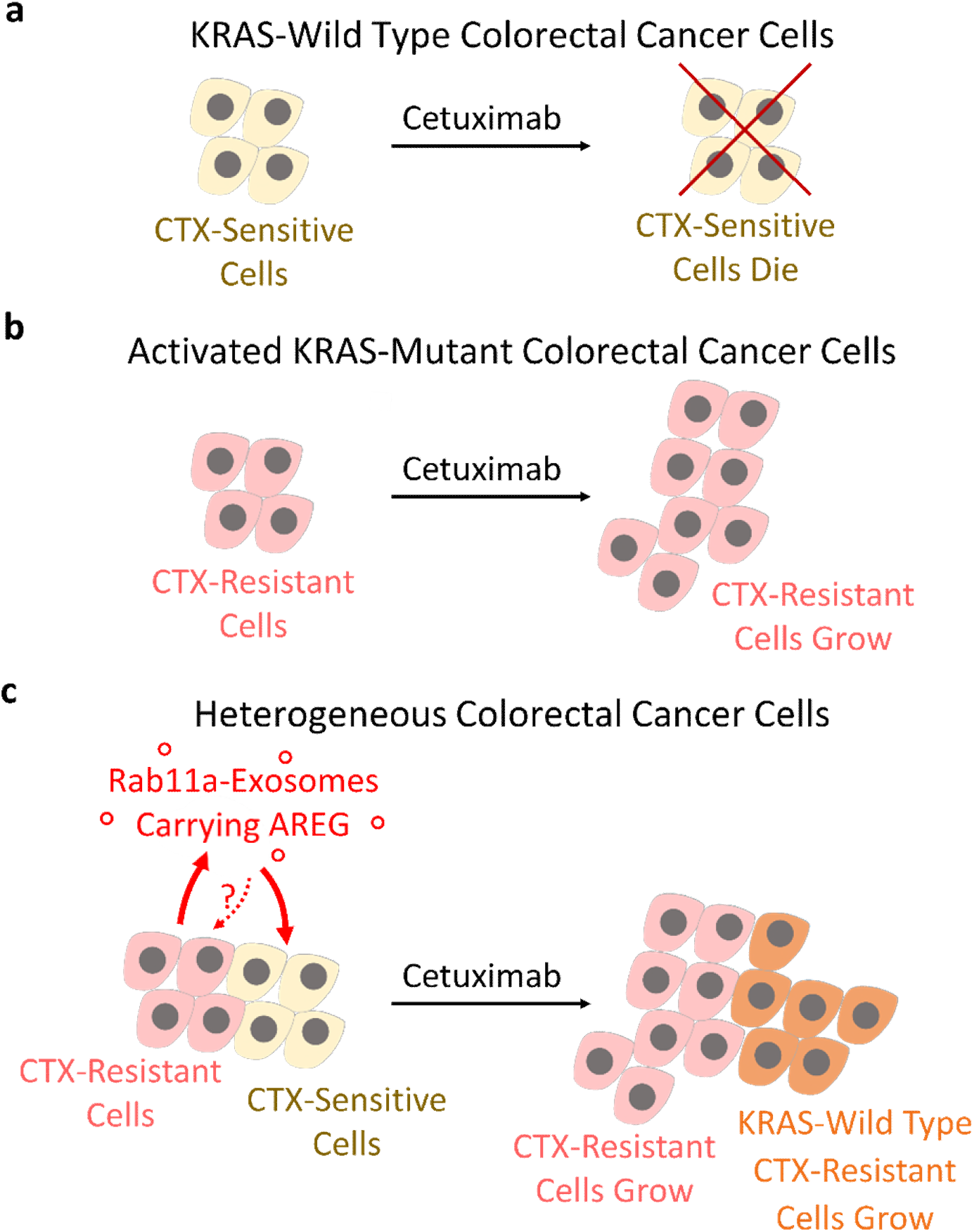
Schematic model highlighting the role of AREG on Rab11a-exosomes in promoting cetuximab-resistance in colorectal cancer. (a) The monoclonal antibody cetuximab (CTX) is typically effective in treating KRAS-wild type colorectal cancer (CRC) cells (yellow cells). (b) Cetuximab is ineffective on other CRC cells that carry an active KRAS mutation (pink cells). These cells may produce Rab11a-exosomes under nutrient and hypoxic stress, but they are presumably not required to maintain cetuximab resistance. (c) In heterogeneous CRC tumours, stress-induced AREG-loaded Rab11a-exosomes from CTX-resistant cells contribute to a mechanism leading to the development of drug resistance in previously CTX-responsive cells, supporting clonal heterogeneity during tumour progression and further evolutionary change (orange cells).

## METHODS

### Cell culture

The CRC cell lines were purchased from ATCC to ensure authenticity. HCT116 cells were cultured in McCoy’s 5A medium (Gibco Life Technologies), Caco-2 cells in EMEM (Gibco Life Technologies) and SW480 and SW620 cells in DMEM GlutaMAX (Gibco Life Technologies). During normal cell culture, all media was supplemented with heat inactivated foetal calf serum (FBS, 10% McCoy’s 5A and DMEM GlutaMAX; 20% EMEM) and 1000 U/ml (1%) penicillin-streptomycin (Gibco Life Technologies). Cells were incubated at 37°C with 5% CO_2_ and were not used beyond passage 20.

### EV isolation

Cells were plated at a density of 9 x 10^6^ cells per plate and cultured until they reached 80% confluency. Following this, the media was changed to serum-free basal medium (DMEM-F12, Gibco Life Technologies) supplemented with 1% insulin-transferrin-selenium solution (ITS, #4140045, Gibco Life Technologies) for 24 hours. For glutamine-depletion isolations, HCT116 cells were grown for 24 hours in DMEM-F12 without L-glutamine (#21331046 Gibco Life Technologies) supplemented with physiological (2 mM) or low (0.15 mM) glutamine (Gibco Life Technologies). The low dose of glutamine was determined by undertaking dose-response experiments with reducing glutamine concentrations to determine a concentration that significantly decreases 4E-BP1 hyperphosphorylation (Fan *et al*, 2020). Torin1 was added at 100 nM for SW480 cells and 150 nM for Caco-2 and SW620 cells.

Following 24 hours of cell growth in the relevant isolation conditions, the medium was harvested at pre-cleared by centrifuging at 500 x g and 2,000 x g both for 10 minutes at 4°C to remove cells, non-vesicular debris and larger vesicles. Following this, the supernatant was filtered using 0.22 µm vacuum filters (Milex). The filtrate was then concentrated using a tangential flow filter with a 100 kDa membrane (Viva-flow 50R, Sartorius), followed by ultrafiltration using 100 kDa Amicon Ultra-15 units (Milex) at 4000xg for 5-10 minutes at 4°C to a final volume of 1 ml. EVs were then isolated using size exclusion chromatography, by injecting the sample onto a 24 x 1 cm size exclusion column containing Sepharose 4B Fast Flow resin (pore size 84 nm) set-up in an AKTA Start System (GE Healthcare Life Science). EVs were eluted in PBS, collecting 30 x 1 ml fractions. Fractions corresponding to the initial EV peak were pooled into 100-kDa Amicon Ultra-4 tubes and centrifuged at 4,000 x g at 4°C to a final volume of approximately 120 µm for analysis. EVs were stored for a maximum of 24 hours at 4°C for transmission electron microscopy or functional analysis, or long term at -80°C for nanoparticle tracker analysis or western blotting.

### Transmission electron microscopy

Approximately 0.2 µg/µl of purified EVs were incubated on glow discharged 300-mesh 3 mm carbon-coated copper grids (TAAB Laboratories) for 2 minutes and then blotted dry and negatively stained with 2% uranyl acetate for 10 seconds. Following this the grid was blotted and allowed to air dry before imaging on a FEI Tecnai T12 transmission electron microscope (FEI UK Ltd) operated at 120 kV with a Gatan Oneview digital camcer (Gatan Inc) at 120,000X magnification.

### Nanoparticle tracker analysis

The NS500 NanoSight^®^ was used to capture five 30 second videos per EV sample (diluted 1:500 in PBS). Particle movement was assessed using NTA software 3.2 (NanoSight Ltd) to generate graphs plotting EV concentration (x 10^8^ ml^-1^) against particle size (nm).

### Western blotting

Cell lysates were collected by lysing cells in RIPA buffer supplemented with protease and phosphatase inhibitors (Sigma) and centrifugation at 14,000 rpm for ten minutes at 4°C. Total protein context was measured using a BCA protein assay kit (Pierce, Thermo Scientific). When undertaking western blotting on EVs, we loaded gels with EV preparations from the same protein mass of secreting cells so that changes in band intensity on the blots with glutamine depletion or Torin1 exposure reflected a net change in secretion of the marker on a per cell basis. We have previously shown that the normalisation method used during western analysis does not affect the overall conclusions made (Fan *et al*, 2020).

Equal amounts of protein or EVs were mixed with reduced (containing 5% β-mercaptoethanol, Sigma) or non-reduced 4 x Laemmli sample buffer (Bio-Rad) and denatured at 95°C for 10 minutes. Samples were then electrophoretically separated on 10% mini-PROTEAN^®^ precast gels (Bio-Rad), followed by wet-transfer onto a Immobilin^®^-Fl polyvinylidene difuoride membrane with a 0.22 µm pore size (Merck) at 100 V for one hour at 4°C. Membranes were then blocked in 5% bovine serum albumin (BSA, Sigma) for reduced conditions or 5% semi-skimmed milk (Sigma) for non-reduced conditions, both in TBS containing 1% Tween-20 (TBST, Sigma) for one hour at room temperature. Membranes where then incubated with primary antibodies diluted in the same blocking buffer overnight at 4°C. The following day, membranes underwent 3 x 10-minute washes in TBST and then probed with the relevant secondary HRP-conjugated antibody. The membranes then underwent a further 3 x 20-minute washes in TBST before incubation in Trident femto Western HRP solution (GeneTex) and visualisation using the ChemiDoc Touch Imaging System (Bio-Rad). Relative band intensities were quantified using ImageJ and normalised to tubulin or cell lysate levels.

Antibody suppliers, catalogue numbers of concentrations used were as follows: rabbit anti-4E-BP1 (Cell Signaling Technology #9644), rabbit anti-S6 (Cell Signaling Technology #2217, 1:4000), rabbit anti-p-S6-Ser240/244 S6 (Cell Signalling Technology #5364, 1:4000), rabbit anti-Akt (Cell Signaling #9272, 1:1000), rabbit anti-p-Akt-Ser473 (Cell Signaling #4060, 1:500), rabbit anti-Caveolin-1 (Cell Signaling Technology #3238, 1:500), goat anti-AREG (R&D Systems #AF262, 1:200), mouse anti-Tubulin (Sigma #T8328, 1:4000), mouse anti-CD81 (Santa Cruz #23962, 1:500), mouse anti-CD63 (BD Biosciences # 556019, 1:500), mouse anti-Alix (Abcam ab117600, 1:500), rabbit anti-Syntenin-1 antibody (Abcam ab133267, 1:500), rabbit anti-Tsg101 (Abcam ab125011, 1:500), mouse anti-Rab11 (BD Biosciences #610657, 1:500), rabbit anti-EGFR (Cell Signaling #4267, 1:1000), rabbit anti-p-EGFR-Tyr1068 (Cell Signaling #3777, 1:500), rabbit anti-p44/42 MAPK (ERK; Cell Signaling Technology #4695, 1:1000), rabbit anti-p-p44/42 MAPK (Cell Signaling Technology #4370, 1:1000), anti-mouse IgG (H+L) HRP conjugate (Promega #W4021, 1:10000), anti-rabbit IgG (H+L) HRP conjugate (Promega #W4011, 1:10000), anti-goat IgG (H+L) HRP conjugate (R&D Systems #HAF109, 1:100).

### Growth assays

Recipient cells were seeded in 96-well plates (HCT116 cells at 2 x 10^3^, SW480 and Caco-2 cells at 3 x 10^3^ and SW620 cells 4 x 10^3^ per well) in 200 µl of medium following pre-treatment with freshly prepared EVs (4 x 10^3^ per cell) for 30 minutes at 37°C. Growth assays were undertaken in reduced serum conditions (HCT116 cells, 1% FBS, SW480, SW620 and Caco-2 cells 0.25% FBS) and measured over five days by live image acquisition using the IncuCyte ZOOM^®^ Live Cell Imager (Essen Bioscience). The IncuCyte analysis software was used to automatically detect cell edges in order to create a confluency mask, which was used to calculate cellular growth and normalised to confluence at time zero, giving a fold change in confluency. Eight technical repeats were employed for each condition and each experiment was repeated at least three times unless otherwise stated.

AREG-neutralising experiments were performed by pre-incubating freshly prepared EVs (at 4 x 10^3^ EVs per recipient cell) with either 4 µg/ml of goat anti-AREG (R&D Systems #AF262) or goat anti-IgG (R&D Systems #AB-1080-C) as a control antibody in the relevant reduced serum media, at 37°C for two hours (Fan et al., 2020). Following this, the EVs were added to the recipient cells and analysed as described above. Experiments investigating EV-free AREG were undertaken by incubating recipient cells with 0.03 – 3000 ng/ml recombinant human Amphiregulin protein (R&D Systems #262-AR-100), for 30 minutes at 37°C before plating as described above.

Drug and inhibitor experiments were undertaken by adding the drug (research grade cetuximab 30 µg/ml, R&D Systems MAB9577) or inhibitor (AG1478 hydrochloride 2.5 mM, Tocris #1276, BMS599626 dihydrochloride 2.5 mM, Tocris #5022) to the recipient cells at the same time as the freshly prepared EVs and incubated for 30 minutes at 37°C before plating.

### Assessment of EV-induced pEGFR and pERK activation

HCT116 or Caco-2 cells were plated in 6-well plates in full FBS supplemented media at a density of 4 x 10^5^ cells per well. After 24 hours the cells were serum starved (0% FBS) overnight and the following day freshly prepared EVs (4 x 10^3^ per cell) were pre-incubated with or without AREG-neutralising antibody (4 µg/ml) for 2 hours at 37°C and added to the recipient cells in fresh 0% FBS media. Cell lysates were then collected after 5 minutes for EGFR analysis and 15 minutes for ERK activation for analysis by Western blotting.

### Statistical analysis

For Western blot analysis, relative signal intensities were analysed using the Kruskal-Wallis or Mann-Whitney U test. For growth assays, data was analysed by two-way Analysis of Variance (ANOVA). All data is presented as mean ± standard deviation and a statistically significant result was defined as p<0.05. Data analysis was performed using GraphPad Prism version 8.3.1 (GraphPad Software, Inc).

## ACKNOWLEDGEMENTS

We are very grateful to Errin Johnson for providing electron micrographs for this study. We acknowledge the support of Cancer Research UK (C19591/A19076, C602/A18974) and the BBSRC (BB/N016300/1, BB/R004862/1). This research was funded in part by the above grants from CRUK and the BBSRC. For the purpose of Open Access, the author has applied a CC BY public copyright licence to any Author Accepted Manuscript (AAM) version arising from this submission.

## Abbreviations

AG1478: EGFR kinase domain inhibitor
AREG: Amphiregulin
ANOVA: Analysis of Variance
BSA: Bovine Serum Albumin
Cav-1: Caveolin-1
CMS: Consensus Molecular Subtype
CRC: Colorectal Cancer
CTX: Cetuximab
dUC: Differential Centrifugation
DMSO: Dimethyl Sulfoxide
EGFR: Epidermal Growth Factor Receptor
ERK: Extracellular Signal-Regulated Kinase 1
ESCRT: Endosomal Sorting Complex Required for Transport
EV: Extracellular Vesicle
FBS: Foetal Bovine Serum
Gln: L-glutamine
HER1: Human EGFR Related 1
HRP: Horse Radish Peroxidase
IgG: Immunoglobulin G
ILVs: Intraluminal Vesicles
ITH: Intratumoral Heterogeneity
ITS: Insulin, Selenium and Transferrin
mTORC1: mechanistic Target of Rapamycin Complex 1
MAPK: Mitogen-Activated Protein Kinase
mCRC: metastatic CRC
MSI: Microsatellite Instability
MSS: Microsatellite Stability
NTA: Nanosight Tracker Analysis
n/a: not applicable
PBS: Phosphate Buffered Saline
SD: Standard Deviation
sEV: small EV
SEC: Size-Exclusion Chromatography
Syn-1: Syntenin-1
TBS: Tris-Buffered Saline
TBST: TBS with 1% Tween-20
TEM: Transmission Electron Microscopy
UCA-1: Uroepithelial Carcinoma-Associated-1

## Supplementary Figures

**Fig. S1.**
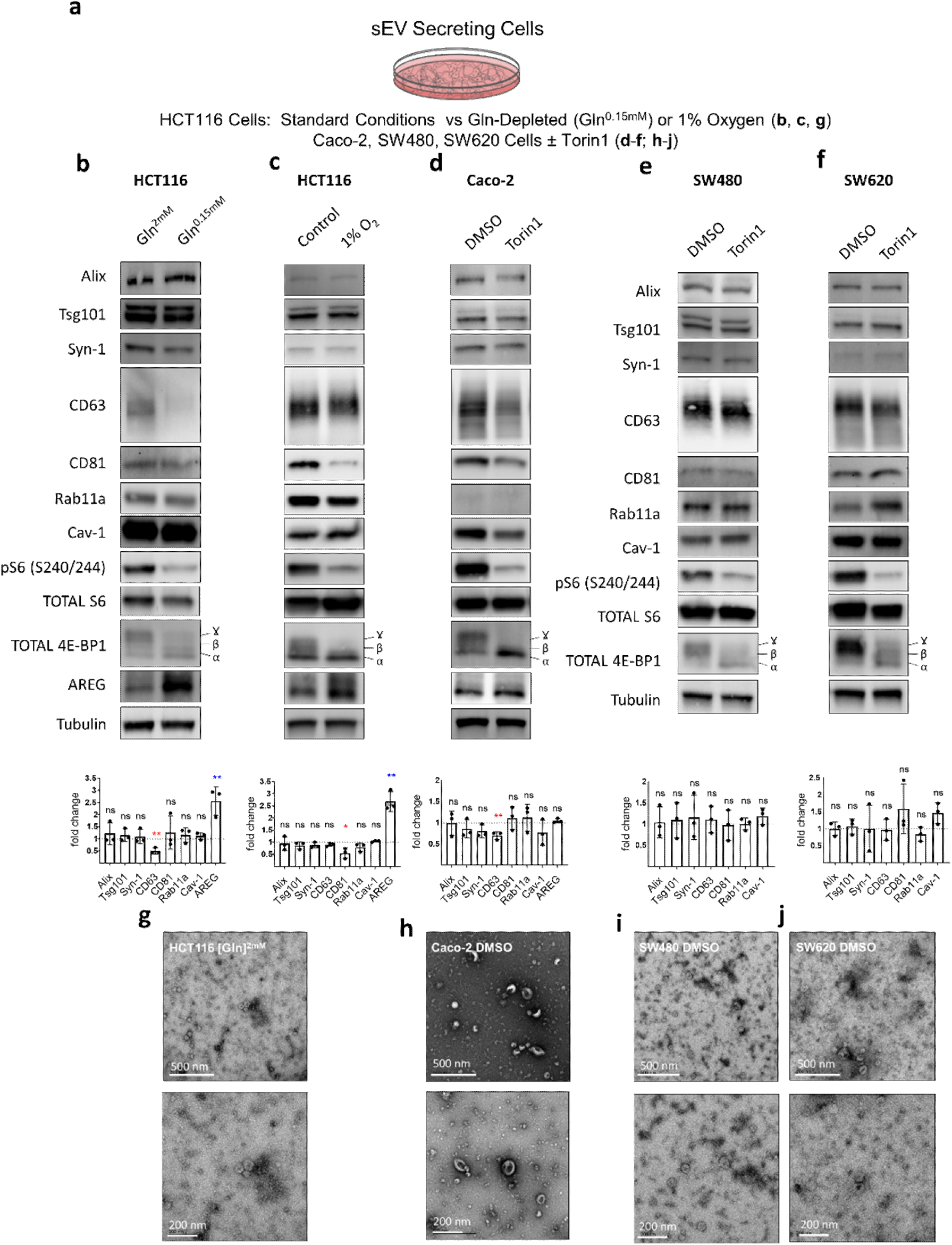
mTORC1 inhibition in CRC cell lines generally does not affect cellular expression levels of Rab11a and other sEV markers. (a) Schematic of manipulations and drug treatments applied to cells and assessed by western analysis in (b-f) below. (b-f) Western blots of CRC cell lysates from cells incubated in mTORC1-inhibitory conditions that induce Rab11a-exosome secretion versus non-inhibitory conditions. The Rab11a-exosome-inducing conditions include glutamine depletion (HCT116 cells, 0.15 mM versus 2 mM; b), hypoxia (HCT116 cells, 1% hypoxia; c), and Torin1 treatment for Caco-2 (150 nM Torin1; d), SW480 (100 nM Torin1; e) and SW620 (150 nM Torin1; f) cells. Note reduced levels of phospho-S6 (P-S6) and phosphorylated forms of 4E-BP1 (α, β, and γ) under inhibitory conditions, as well as similar levels of many sEV markers (Alix, Tsg101, Syn-1, CD81, Rab11a, Cav-1) following most treatments, as shown in histograms (levels normalised to Tubulin; n = 3). CD63 expression is reduced when mTORC1 is inhibited in glutamine-depleted, but not hypoxic, HCT116 cells and in Caco-2 cells. (g-j) Transmission electron microscopy images of sEV preparations confirm that vesicles isolated from these different cells are less than 200 nm in diameter and reveal a classic cup-like morphology of vesicles prepared using this procedure. *P<0.05. Red and blue asterisks denote reduction and increase in protein levels respectively in mTORC1-inhibited cells.

**Fig. S2.**
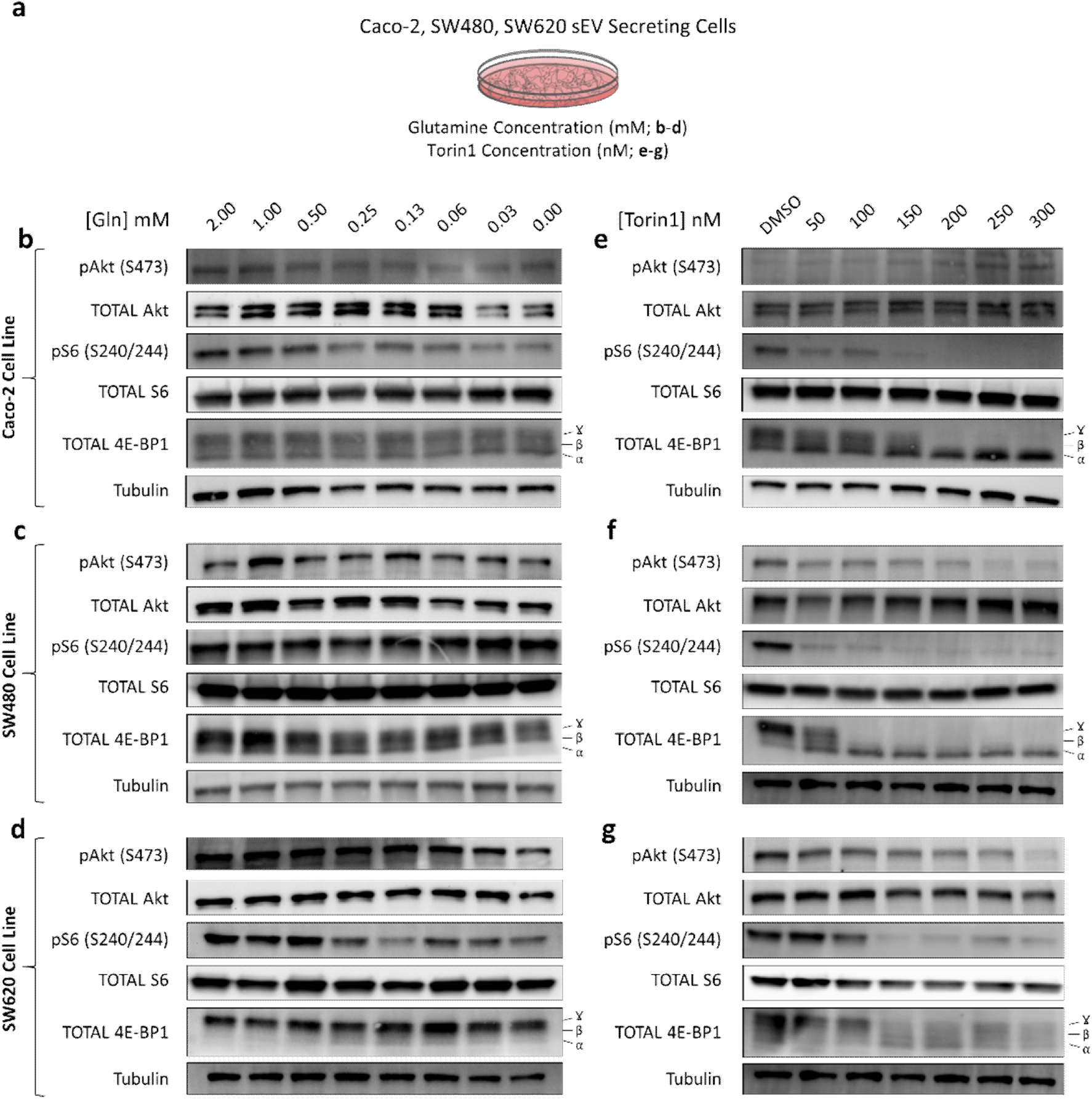
Torin1, but not glutamine depletion, inhibits mTORC1 activity in Caco-2, SW480 and SW620 cells. (a) Experimental design relevant to the data below. (b-d) Western blots of cell lysates from Caco-2 (b), SW480 (c) and SW620 (d) cells cultured in different concentrations of glutamine reveal that the activity of Akt, and mTORC1 downstream targets, S6 and 4E-BP1, as measured by their phosphorylation, is largely unaffected by glutamine depletion. Phosphorylated forms of 4E-BP1 run slower in polyacrylamide gels, producing α, β, and γ forms. (e-g) Western blots of cell lysates from Caco-2 (e), SW480 (f) and SW620 (g) cells cultured in different concentrations of mTOR inhibitor Torin1, which inhibits mTOR-containing mTORC2 (a regulator of Akt) as well as mTORC1. These reveal that the activity of Akt, and mTORC1 downstream targets, S6 and 4E-BP1, as measured by their phosphorylation, is blocked by Torin1 in a dose-dependent manner.

**Fig. S3.**
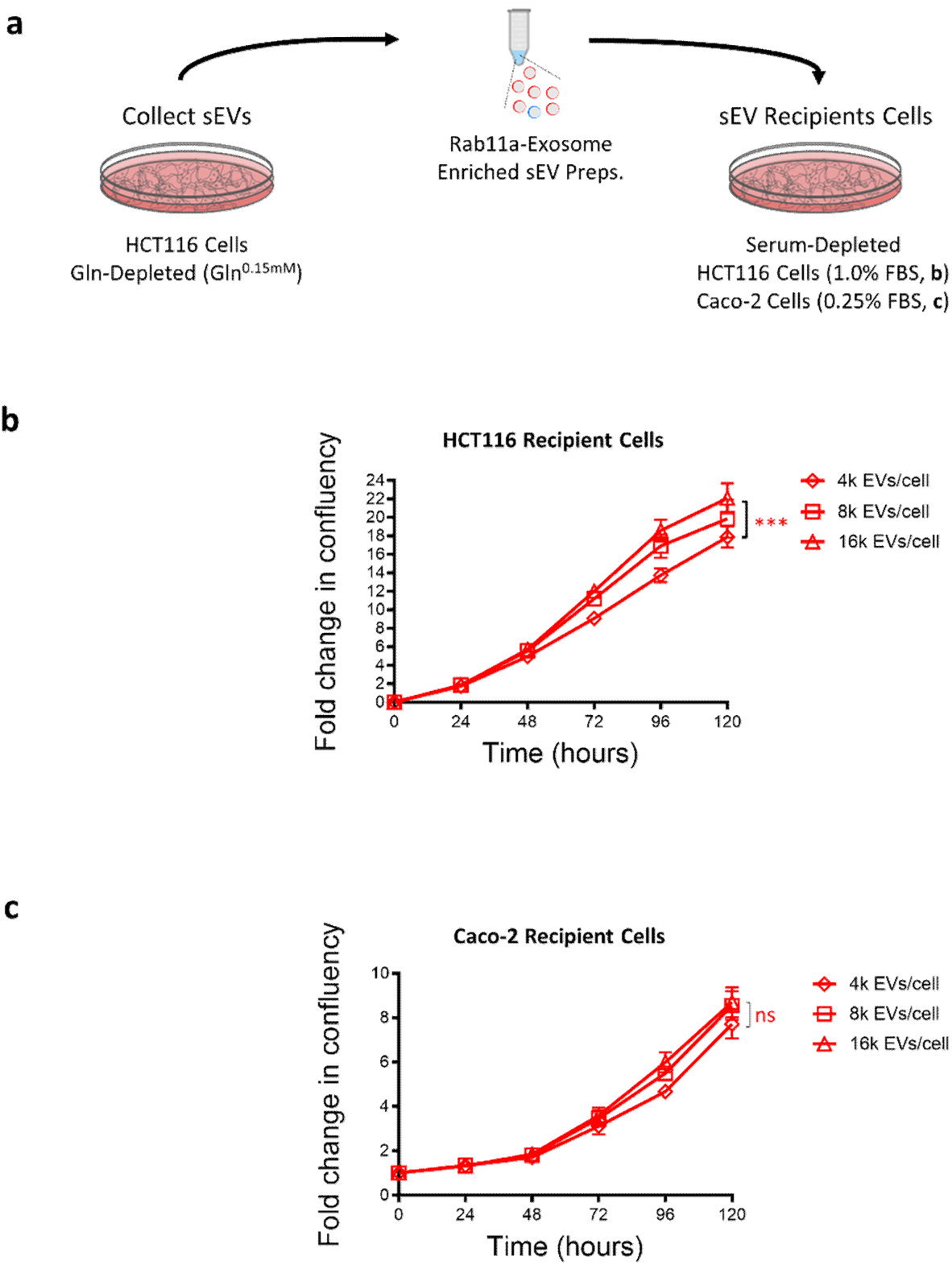
Rab11a-exosome preparations from HCT116 cells promote growth of HCT116 cells in a dose-dependent fashion under serum-depleted conditions. (a) Experimental design relevant to the data below. (b) Growth-promoting activity of HCT116 Rab11a-exosome-enriched sEV preparations increases with dose for naïve HCT116 cells under 1% FBS conditions. (c) Growth-promoting activity of HCT116 Rab11a-exosome-enriched sEV preparations does not significantly increase with dose above 4000 sEVs per target cell for naïve Caco-2 cells under 0.25% FBS conditions. Eight technical repeats were employed for each condition. ***P<0.001.

**Fig. S4.**
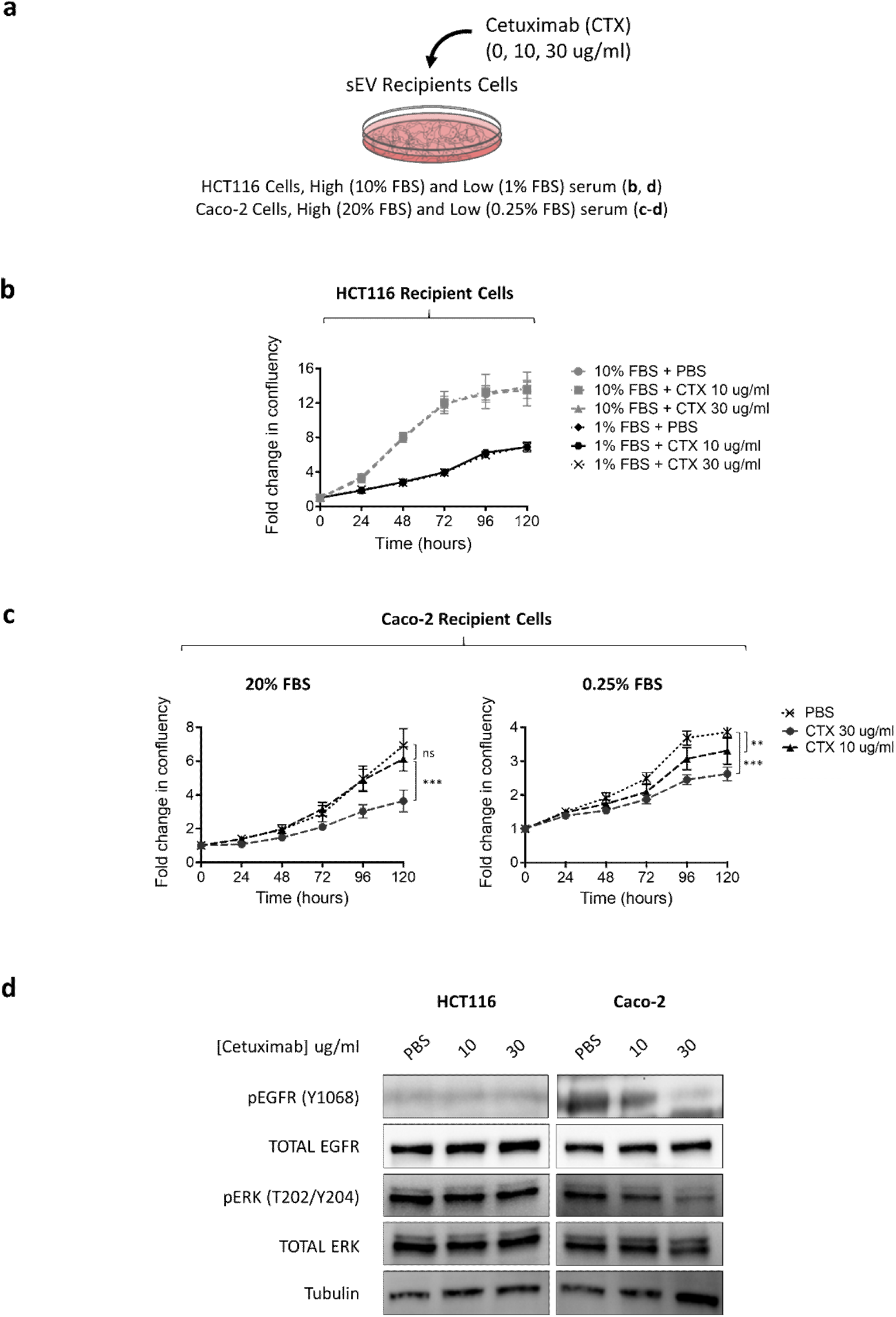
Cetuximab only reduces the growth of CRC cells carrying wild type KRAS. (a) Experimental design relevant to the data below. (b) Cetuximab at 10 and 30 µg/ml concentration has no effect on HCT116 cells grown under either high (10% FBS) or low (1%) serum conditions. Cetuximab resistance has previously been shown to involve the activated KRAS mutation in CRC cells. (c) Cetuximab inhibits growth of Caco-2 cells, which possess wild type KRAS, under high (20%) and low (0.25%) serum conditions in a dose-dependent manner. (d) Western blot of cell lysates showing that cetuximab treatment has no effect on EGFR or ERK phosphorylation in HCT116 cells, but reduces the phosphorylation of both proteins in Caco-2 cells in a dose-dependent manner. In growth experiments, eight technical repeats were employed for each condition and each experiment was repeated three times. **P<0.01; ***P<0.001.

**Fig. S5.**
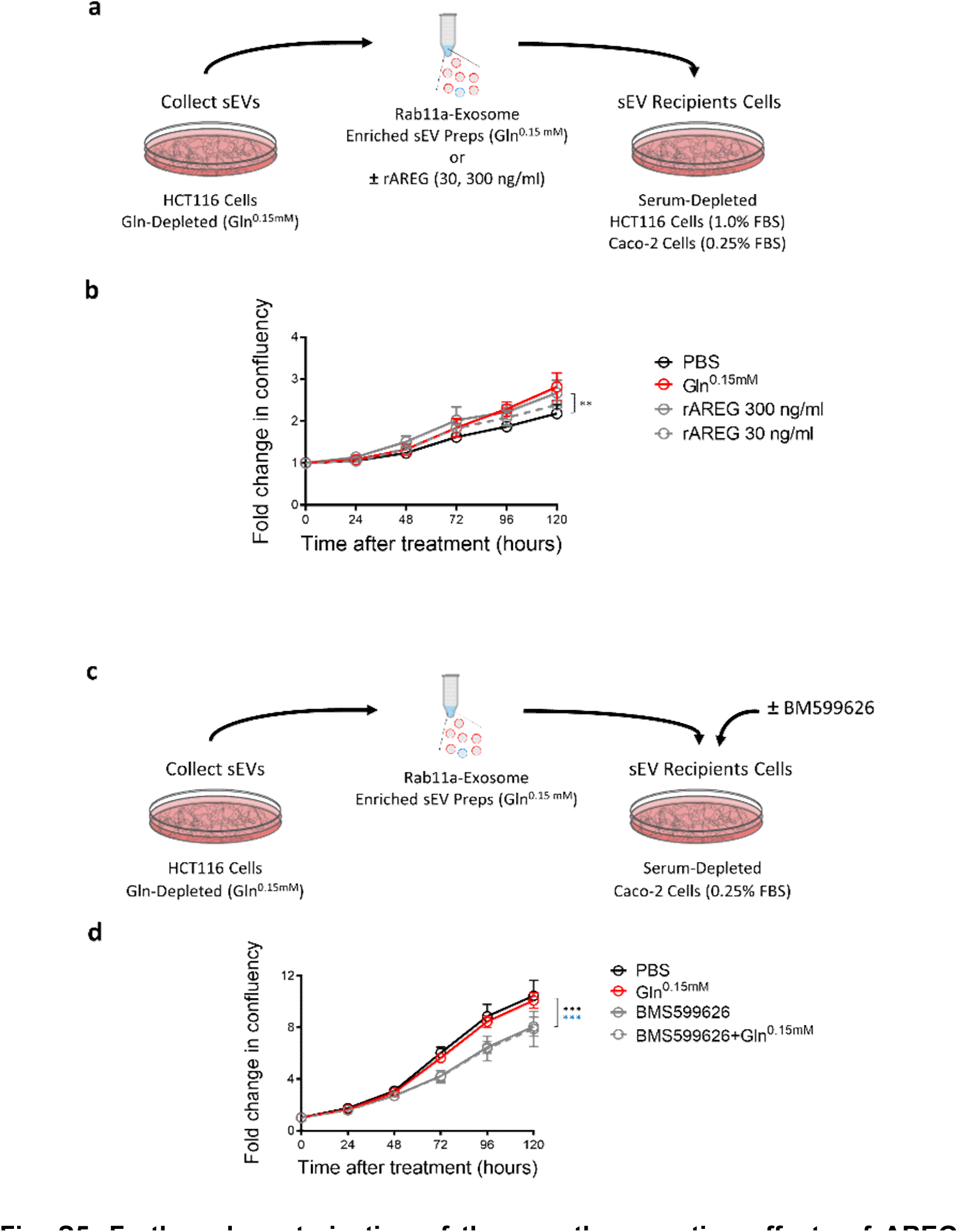
Further characterisation of the growth promoting effects of AREG on Rab11-exosomes. (a) Experimental design relevant to (b). (b) Comparison of the growth-promoting effects on HCT116 cell growth over a 120-hour time course using soluble recombinant AREG (rAREG) and AREG on Rab11a-exosome-enriched sEV preparations (Gln^0.15 mM^). Black asterisks denote significant difference between 300 ng/ml rAREG-induced growth and control. (c) Experimental design relevant to (d). Schematic of experimental approach employed used to test the effect of BMS599626. (d) Comparison of the effect on Caco-2 cell growth over a 120-hour time course of Rab11a-exosome-enriched sEV preparations and PBS control in the absence and presence of pan-HER receptor tyrosine kinase inhibitor BMS599626 inhibitor. This experiment was repeated twice. For growth curves, eight technical repeats were employed for each condition. **P<0.01; ***P<0.001. In (d), black and blue asterisks denote significant inhibitory effects on PBS control and Rab11a-exosome-treated cells respectively.

